# Regional Variations in Live Proportions of Southwest Pacific Cold-Water Coral *Solenosmilia variabilis* Reefs

**DOI:** 10.1101/2024.04.08.588521

**Authors:** Kelsey Archer Barnhill, Caroline Chin, Di Tracey, Malcolm Clark, Laurence H. De Clippele, Sian F. Henley, Uwe Wolfram, Sebastian Hennige

**Affiliations:** School of GeoSciences, University of Edinburgh, Edinburgh, UK; National Institute of Water and Atmospheric Research, Wellington, New Zealand; School of Biodiversity, One Health & Veterinarian Medicine, University of Glasgow, Glasgow, UK; Technische Universität Clausthal, Clausthal-Zellerfeld, Germany

**Keywords:** *Cold-Water Corals*, *Deep-Sea*, *Framework-Forming Cold-Water Corals*, *Graveyard Seamounts*, *Louisville Seamount Chain*, *New Zealand*, *Reef-Building Cold-Water Corals*, *Seamount*, *Solenosmilia variabilis*

## Abstract

Reef-building cold-water-corals (CWC) form deep-sea habitats that can create biodiversity hotspots. As live coral and dead intact framework provide disparate ecosystem services and are vulnerable to different anthropogenic stressors, it is important to quantify the proportions of each on CWC reefs. We analysed 1,160 images of *Solenosmilia variabilis* reefs at four sites off New Zealand (Valerie and Forde Guyots at the Louisville Seamount Chain & Ghoul and Gothic Seamounts at the Graveyard Seamount Complex) to determine the ratio of live coral to the whole reef area (termed live:reef). We found live:reef ratios are significantly different between sites at the offshore Louisville Seamount Chain and onshore Graveyard Seamount Complex. This could be driven by reef position relative to the aragonite saturation horizon (ASH) as corals in the Louisville Seamount Chain live below the ASH in colder and deeper waters than those at the Graveyard Seamount Complex, which live above the ASH. In the southwest Pacific, depth is a driver of live:reef ratios, with a larger proportion of live coral at shallow depths and dead intact framework at deeper depths. The live:reef ratios at Gothic Seamount within the Graveyard Seamount Complex remained stable between 2015 and 2020 despite significant differences in live coral, dead intact framework, and reef structure surface area. Our results indicate live:reef ratios can be used to estimate the amount of dead intact framework threatened by shoaling ASH due to ocean acidification at each site, which can help inform which sites could be protected as possible climate change refugia.

## Introduction

Scleractinian reef-building cold-water corals (CWC) have the ability to form deep-sea habitats, which can create biodiversity hotspots (D’Onghia et al., 2010; Henderson et al., 2020; Roberts et al., 2006; Thresher et al., 2014). Live, tissue-covered colonies, of framework-forming corals can be found growing atop a 3-dimensional structurally complex skeleton that is exposed to seawater (referred to throughout this study as ‘dead intact framework’) and increase local biodiversity by providing a variety of habitat niches (Mortensen et al., 1995; Price et al., 2019; Schnabel et al., 2019) (Figure 1). When a coral colony reaches a certain size, parts of the colony become sheltered from the currents and therefore food supply causing those parts to die off (Corbera et al., 2022; Georgoulas et al., 2023; Hennige et al., 2021; Mienis et al., 2007); over time the dead intact framework will make up the majority of a colony (Vad et al., 2017) and eventually a reef (Mortensen et al., 1995). It is the dead intact framework that has the highest associated biodiversity and contributes most to carbon and nitrogen cycling (Buhl-Mortensen et al., 2010; De Clippele et al., 2021; Henry et al., 2010; Henry & Roberts, 2007). The largest quantity of CWC-supported biodiversity is found in the reef transition zone, i.e., the zone where there is a large proportion of dead intact framework (Mortensen & Fosså, 2006). Different species of fish use CWC reef habitats for various purposes, including shelter (Hebbeln et al., 2009), feeding (D’Onghia et al., 2012), spawning and nurseries (D’Onghia et al., 2010; Henry et al., 2013, 2016). Just as the reef can be used by animals in different ways, some interact more with the live corals or dead intact framework (Kazanidis et al., 2021; Mortensen & Fosså, 2006;), an example being fish at Porcupine Bank in the NE Atlantic, where the morid cod *Guttigadus latifrons* was mostly present among living corals while the oreo *Neocyttus helgae* was mostly present among the dead intact framework (Söffker et al., 2011).

**Figure 1.**
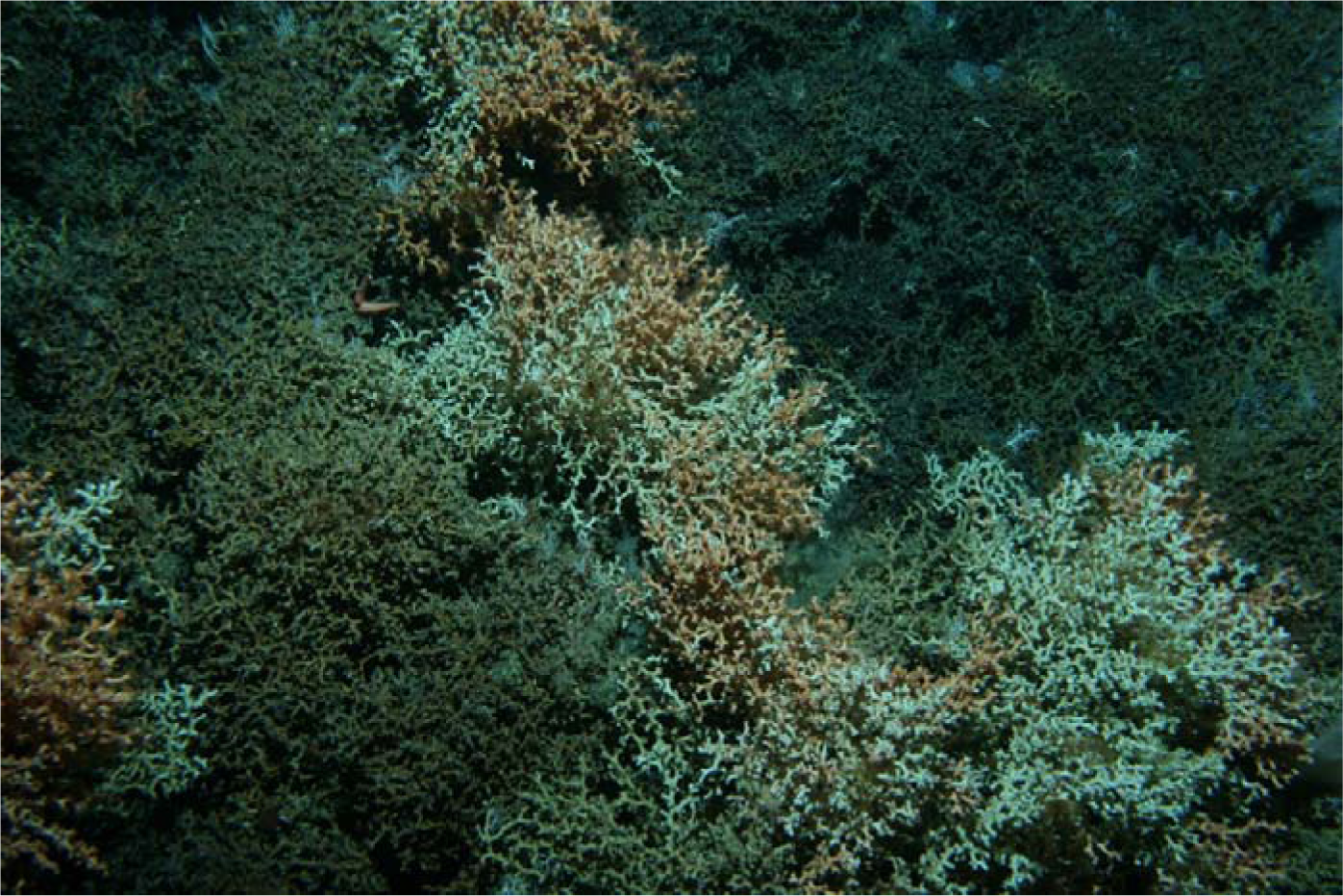
Image of the Scleractinian reef-building CWC *Solenosmilia variabilis* from Gothic Seamount at the Chatham Rise. The orange and white sections of coral are the youngest parts of the reef, representing live colonies while the brown reef material is dead intact framework. Photo: Deep-Towed Imaging System, NIWA.

Globally, CWCs primarily exist in waters saturated with respect to aragonite, with occurrences largely constrained by the aragonite saturation horizon (ASH) (Ω_ARAG_ > 1) (Davies & Guinotte, 2011), which ranges in depth from 200-1,320 m in the southwest Pacific Ocean (Feely et al., 2002). Aragonite saturation state is largely determined by the amount of atmospheric CO_2_ taken up by the ocean, where greater CO_2_ uptake drives lower Ω_ARAG_ (Feely et al., 2004). However, unlike other framework-forming species, *Solenosmilia variabilis* (Duncan, 1873) inhabits waters of a lower aragonite saturation state at many locations, such as on seamounts in Australia’s Huon Marine Reserve and Tasman Fracture Zones, where it is found along or below the ASH from Ω_ARAG_ = 0.83-1.1 (Thresher et al., 2011). *S. variabilis* is globally distributed, and, in the absence of *Desmophyllum pertusum* (synonymised from *Lophelia pertusa* (Addamo et al., 2016)), is the key framework-forming CWC species in the southwest Pacific Ocean, representing the largest distribution and biomass of reef-building CWCs in the region (Williams et al., 2020). In the southwest Pacific, 85% of *S. variabilis* occurrences are reported between 800-1,400 m, with 1,000 m as the mean depth (Bostock et al., 2015; Tracey et al., 2013), placing it deeper than other scleractinian framework-forming coral species. Additionally, southwest Pacific *S. variabilis* generally grows in temperatures 3-5℃ cooler than framework-forming corals in the North Atlantic Ocean (Fallon et al., 2014).

Within New Zealand’s extended continental shelf, 62% of *S. variabilis* observations occur on seamounts despite being distributed generally across rises, ridges, and slopes in the region (Tracey, Rowden, et al., 2011). Mean biomass of *S. variabilis* has been found to be 29 times higher on seamounts (xL = ∼347.17 g·wet weight (ww)·m^-2^) than at comparable depths on the nearby continental slope (xL = ∼11.94 g·ww·m^-2^) (Rowden et al., 2010). On New Zealand southwest Pacific seamounts, *S. variabilis* dominates, contributing 73% of all biomass assemblages (Rowden et al., 2010). Seamounts are habitats for abundant populations of demersal fish species (Rogers, 2018), such as the orange roughy (*Hoplostethus atlanticus*), which has been commercially fished in New Zealand since the 1980s and was once the country’s most economically valuable fish species (Clark, 1995). These increased levels of fish density led to seamounts being among the most heavily fished features in New Zealand, causing an overlap in trawling areas and CWC presence (Clark, Bowden, et al., 2010; Clark et al., 2009, 2019; Clark & Koslow, 2008).

Several southwest Pacific fisheries, including the orange roughy, have recorded *S. variabilis* bycatch at sites such as Chatham Rise and Campbell Plateau (Anderson & Clark, 2003; Tracey, Baird, et al., 2011). At sites farther away from the continental slope, such as the Louisville Seamount Chain, seamounts comprise all suitable orange roughy habitat as there is no adjacent slope, making them key locations for trawling (Clark, Dunn, et al., 2010). Reef habitats that have been heavily trawled, are flattened and therefore lose their habitat complexity. They then become unable to provide the same level of ecosystem services as untrawled reefs (Tracey & Hjorvarsdottir, 2019). Using seafloor image data from New Zealand’s Chatham Rise, Clark et al. (2010) suggests just ten trawls on a seamount can reduce coral presence from ∼15-20% cover to zero. Trawling is a major anthropogenic threat to seamount CWC communities, as little evidence of recovery was found 15 years after trawling closure at Chatham Rise, indicating decade-length timescales of recovery (Clark et al., 2019).

The habitat created by *S. variabilis*, along with other stony corals, is classified as a vulnerable marine ecosystem (VME) (FAO, 2009) and is among the ten indicator taxa (species that reflect a site’s environmental state, biodiversity and environmental change) identified by the South Pacific Regional Fisheries Management Organisation for VMEs (Parker et al., 2009). At Australian and New Zealand seamounts, *S. variabilis* is the primary species that forms coral thickets (Williams et al., 2010). In addition to forming habitat, the structural complexity of the features built by this species is also important for early life stages of deep-sea chondrichthyans. Catshark egg cases (genus *Apristurus*) attached atop *S. variabilis* reef structures were found at 1,000 m depth at Gothic Seamount (Armstrong, 2022). In New Zealand, *S. variabilis* along with most coral species, is protected by the Wildlife Act of 1953 and its 2010 amendment (Tracey & Hjorvarsdottir, 2019). Under the criteria described by Townsend et al. (2008), *S. variabilis* in New Zealand is classified as ‘At Risk’ due to declining numbers (Freeman et al., 2013).

Due to slow growth rates (Fallon et al., 2014) and low genetic variation (Zeng et al., 2016), *S. variabilis* is vulnerable to anthropogenic disturbances. Off NE Tasmania, *S. variabilis* reef mounds slowly accumulate to reach 2-3 m in height (Fallon et al., 2014). This reef-forming species is slow growing with colony growth rates ranging from 0.84 -1.25 mm linear extension per year at Tasmanian seamounts, with higher growth rates found at deeper sites (Fallon et al., 2014). At this rate, it would take ∼2,000 years for a *S. variabilis* colony to reach a height of 1 m (Tracey et al., 2023; Tracey & Hjorvarsdottir, 2019). Their growth and accumulation rates are significantly lower than other scleractinian reef-forming species such as *D. pertusum* and *Oculina vericosa*, likely due to species-specific differences or discrepancies in regional productivity levels (Tracey & Hjorvarsdottir, 2019). *S. variabilis* reef habitats persist on scales of thousands of years in the deep sea, with one southwest Pacific fragment having been aged up to ∼47,400 years BP (Fallon et al., 2014).

In the southwest Pacific, this species is found in isolated populations due to the selection of asexual reproduction over larval dispersal, causing limited ecological connectivity between seamounts (Miller & Gunasekera, 2017). *S. variabilis* from Louisville Seamount Chain, located NE of New Zealand in international waters, is placed in a different genetic cluster than samples from Chatham Rise, located within New Zealand’s exclusive economic zone (EEZ) likely caused by a gene flow barrier isolating populations on the two features (Zeng et al., 2017). Similarities in the genetic results obtained on two samples of *S. variabilis* collected from Valerie Guyot in 2014 suggest low genetic variation in the species’ mitochondrial genome (Zeng et al., 2016).

### Study Sites

The Louisville Seamount Chain is a 4,000 km long volcanic hotspot feature with over 80 seamounts that lies in international waters within the South Pacific Regional Fisheries Management Organisation Convention Area, off the eastern coast of NZ (Rowden et al., 2017). Within the seamount chain, there are many seamount features with a flat top due to erosion, called guyots. Within the Louisville Seamount Chain, *S. variabilis* forms reefs ranging from 600 m^2^ to 0.04 km^2^ (Rowden et al., 2017). For this study, we selected Valerie and Forde Guyots as study sites for this region (Figure 2). As of 2017, Valerie Guyot had been trawled 1,826 times and remains open to bottom trawling while Forde Guyot has been trawled 370 times and was closed to bottom trawling in 2005 (Rowden et al., 2017). Valerie Guyot’s summit peaks at 770 m depth with steep flanks from 1,100 m (Clark et al., 2014). Forde Guyot’s summit is a large plateau between 1,000-1,100 m deep with steep flanks from 1,150 m (Clark et al., 2014). Valerie is the larger of the two guyots with an area of 678 km^2^ whilst Forde’s area is 253 km^2^ (Rowden et al., 2017). Both features have portions of their flanks below the ASH (Bostock et al., 2015).

**Figure 2.**
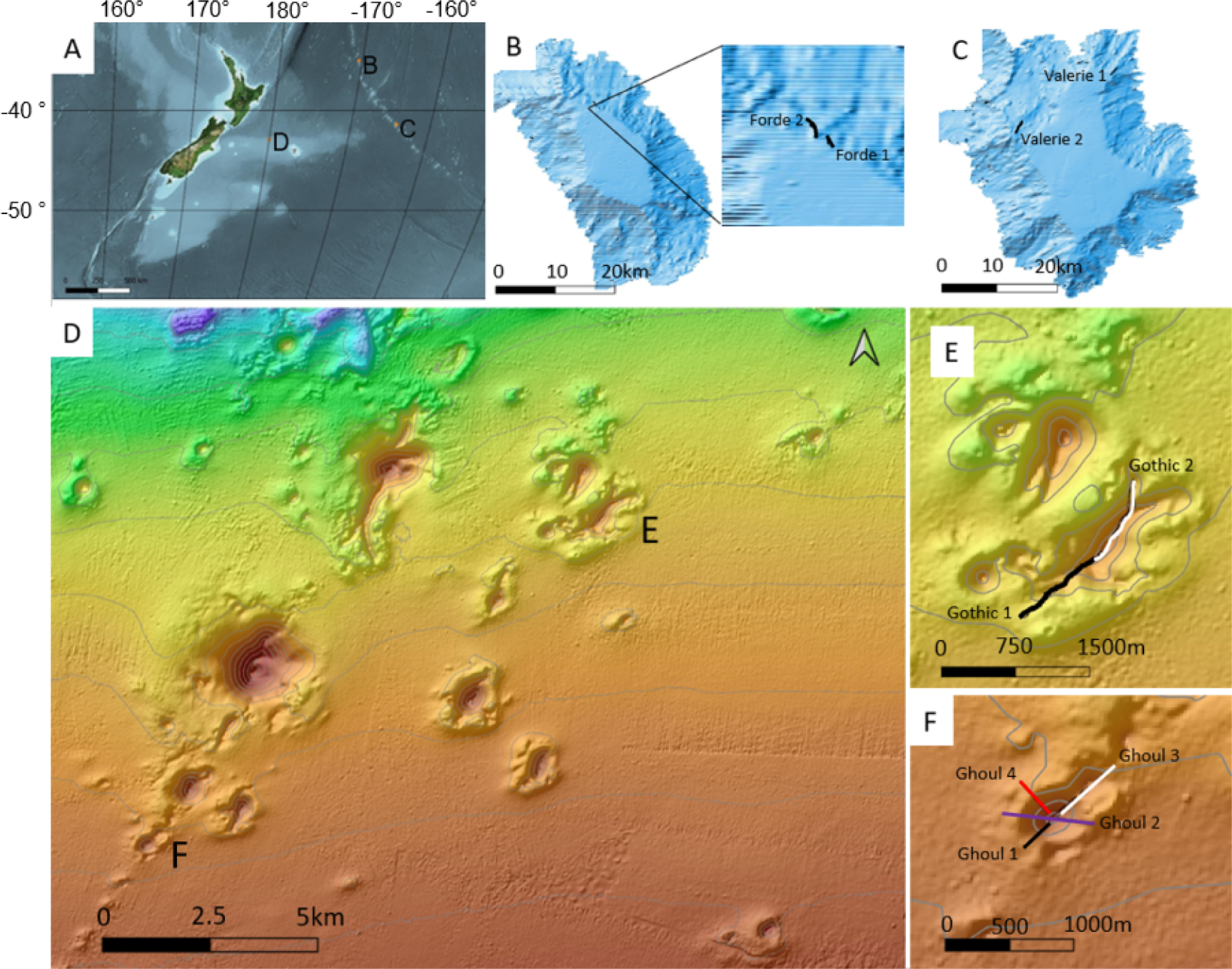
Map of the region with studied features enlarged. A. Map of region with Forde Guyot (B), Valerie Guyot (C) and the Graveyard Seamount Complex (D), denoted. B. Forde Guyot on the Louisville Seamount Ridge with the stations Forde 1 and Forde 2. C. Valerie Guyot on the Louisville Seamount Ridge with the stations Valerie 1 and Valerie 2. D. The Graveyard Seamount Complex within Chatham Rise with Gothic (E) and Ghoul (F) Seamounts denoted. E. Gothic Seamount with the stations Gothic 1 and Gothic 2. F. Ghoul Seamount with the stations Ghoul 1, Ghoul 2, Ghoul 3, and Ghoul 4.

The Graveyard Seamount Complex lies within New Zealand’s EEZ and consists of 28 continental mid-plate volcanic cones close to the northern flank of Chatham Rise (Clark et al., 2006), one of the country’s most productive fishing grounds (McGregor et al., 2019). Within the Graveyard Seamount Complex, *S. variabilis* forms reefs ranging from 100 m^2^ to 0.05 km^2^ (Clark et al., 2021; Tracey et al., 2023). For this study, we selected Ghoul and Gothic Seamounts as study sites for this region (Figure 2). Despite heavy trawling at the seamount complex, Ghoul Seamount has never been trawled and Gothic Seamount has not been trawled since four recorded attempts prior to 2001 (Clark et al., 2019), when it was closed to bottom trawling (Brodie & Clark, 2003). Ghoul Seamount’s depth range from peak to base is 935-1,050 m and Gothic has a slightly larger elevation range of 987-1,160 m (Rowden et al., 2002). Gothic is the larger of the two seamounts with an area of 2 km^2^ whilst Ghoul’s area is 0.6 km^2^ (Rowden et al., 2002). Both features lie entirely above the ASH (Bostock et al., 2015).

### Motivations and Aims

Resilience of fragile benthic communities, such as reef-building scleractinian corals, are defined in part by growth patterns (Hughes, 1987). Assessing CWC growth characteristics, which are driven by environmental conditions and coral biology, is a simple way to quantify coral health in remote locations (Vad et al., 2017). At CWC reefs, live corals and dead intact framework are threatened by different climate change stressors (Barnhill et al., 2023). For example, live CWC colonies are vulnerable to increased temperatures (Brooke et al., 2013), deoxygenation (Dodds et al., 2007) and decreased food supplies (da Costa Portilho-Ramos et al., 2022), whilst the dead intact framework is vulnerable to dissolution under conditions of decreased pH, threatening the structural integrity of reef formations (Hennige et al., 2020; Wolfram et al., 2022). As live coral and dead intact framework provide disparate ecosystem services and are vulnerable to different anthropogenic stressors, it is important to quantify the proportions of each on CWC reefs.

Assessing the proportion of live corals on a CWC reef (hereafter referred to as live:reef ratios) during field surveys has been suggested as a parameter to monitor long-term framework-forming CWC reef health (Vad et al., 2017). Live:reef ratios have previously been explored for the reef-forming species *D. pertusum* (Vad et al., 2017) and *Madrepora oculata* (Orejas et al., 2021). Live:reef ratios have not been determined for the species *Solenosmilia variabilis*. The growth patterns for both *D. pertusum* and *M. oculata* indicate that the proportion of live corals is almost always smaller than the proportion of dead intact framework (Orejas et al., 2021; Vad et al., 2017). In *D. pertusum* colonies, a negative correlation has been found between whole colony size and the live:reef ratio, suggesting the proportion of live polyps decrease as the colony’s growth rate slows down (Vad et al., 2017).

In this study, the objectives are to: (1) explore live:reef ratios between and within four *S. variabilis* reef sites off the coast of New Zealand to understand spatial differences in ratios; compare live:reef ratios between 2015 and 2020 at the repeatedly surveyed Gothic Seamount within the Graveyard Seamount Complex to explore ratio stability over time; and explore the impact a reef’s position in relation to depth and the ASH has on live:reef ratios. These results will be integrated into a discussion of potential environmental drivers influencing live:reef ratios.

## Methods

### Videos & Image Processing

Video footage, collected during three research expeditions on the R/V *Tangaroa* was used for this study: TAN1402 at the Louisville Seamount Chain (January – March 2014), TAN1503 at the Graveyard Seamount Complex (March-April 2015), and TAN2009 (August 2020) at the Graveyard Seamount Complex. All data were collected using the National Institute of Water and Atmospheric Research’s (NIWA) towed camera, attached to the Deep-Towed Imaging System (DTIS). At the Louisville Seamount Chain, we selected two stations from Valerie Guyot (Valerie 1 & 2) and two stations from Forde Guyot (Forde 1 & 2). At the Graveyard Seamount complex, we selected four stations from Ghoul Seamount (Ghoul 1, 2, 3 & 4) and two stations at Gothic Seamount first visited in 2015 (Gothic 1 2015 & Gothic 2 2015) and again in 2020 (Gothic 1 2020 & Gothic 2 2020). Video footage from each station equates to a photographic image transect covering a distance of ∼0.2-0.6 km. The DTIS is a piece of deep-sea sampling equipment that records high-definition video footage (Sony HD 1080i format) from 2-3 m above the seafloor whilst travelling at 0.5-1 knots (Bowden & Jones, 2016; Clark et al., 2019; Hill, 2009; Rowden et al., 2017). The DTIS uses an ultra-short baseline positioning system (USBL, Kongsberg HiPAP) to record seabed position within 1 m accuracy (Rowden et al., 2017). Altitude above the seabed and DTIS depth are continually recorded.

Image frames were extracted from every 10 seconds of each ∼1 hr transect of video footage to create layers in Adobe Photoshop Release 24.1.0. Scaling lasers were annotated and the entire portion of live corals and dead intact framework was labelled in each image following the method of van der Kaaden & De Clippele (2021) and De Clippele et al. (2021) (Figure 3). Blurry, overlapping, or images without coral or scaling lasers in the image were excluded from analysis. In total, 1,160 images were annotated and exported as tiff files for analysis: 117 from Valerie Guyot (Valerie 1: 50; Valerie 2: 67), 64 from Forde Guyot (Forde 1: 11; Forde 2: 53), 273 from Ghoul Seamount (Ghoul 1: 61; Ghoul 2: 60; Ghoul 3: 81; Ghoul 4: 73), and 706 from Gothic Seamount (Gothic 1 2015: 224; Gothic 2 2015: 211; Gothic 1 2020: 158; Gothic 2 2020: 127) (Figure 2).

**Figure 3.**
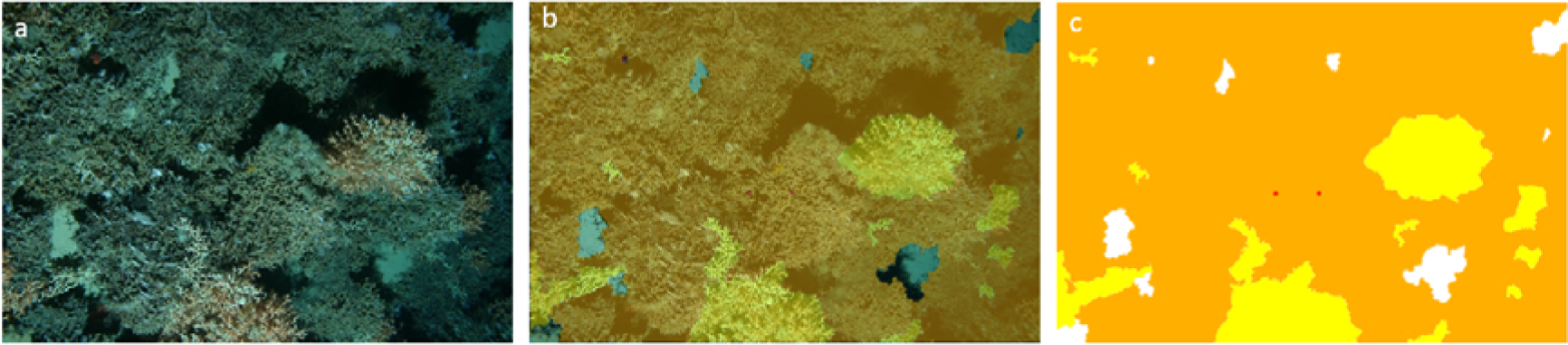
Semi-automatic image annotation process. A. Seafloor image extracted from DTIS video. B. Live corals and dead intact framework highlighted atop the image along with scaling lasers positioned 20 cm apart. C. Layer with manual annotations exported for analysis. Live corals are represented as yellow and dead intact framework is represented as orange.

### Data Analysis

All analyses were completed in RStudio 2022.02.2 Build 485 ‘Prairie Trillium’ Release (Rstudio Team, 2019) using R version 4.2.0 (R Core Team, 2022) and plots were created using the R package ggplot2 version 3.4.2 (Wickham, 2016). We used existing open-source R code stored in Zenodo to calculate total surface area and percentage of live and dead reef portions based on the labelled sections in each image (Figure 3) (van der Kaaden & De Clippele, 2021). From these data, we determined live:reef ratios in each image. We tested the live:reef data for normality using the Shapiro-Wilk Test (Shapiro & Wilk, 1965). Since the data were not normally distributed, we used the non-parametric Kruskal-Wallis rank sum & Wilcoxon rank sum tests to compare differences between and within sites (Kruskal & Wallis, 1952; Wilcoxon, 1945). Using USBL data from the DTIS, we matched 917 of the images with their depth records. Non-parametric linear regressions were used to explore the relationship between water depth and live:reef ratios. Summary data from CTD casts during the TAN1402 expeditions were provided by Owen Anderson of NIWA. Water property variables beyond depth and aragonite saturation were not available at a high enough resolution to differentiate between the four sites, and thus are not included in the analysis. However, depth can act as a proxy for variables such as temperature, salinity, and surface-derived production (Rowden et al., 2017).

## Results

Transect depths at the Louisville stations ranged from 1,081-1,584 m while Graveyard stations were shallower, ranging from 923-1,166 m. These deeper depths led to lower temperatures at the Louisville stations (3.3-3.9°C (Clark et al., 2015; Gammon et al., 2018) compared to 5.2-6.1°C (Goode et al., 2021) at the Graveyard Complex) and aragonite saturation states at or below Ω_ARAG_ = 1 (Anderson, unpublished data). The DTIS surveys at the Graveyard Seamount Complex occurred at depths above the 1,300 m ASH, while those at the Louisville Seamount Chain occurred at depths at or below the 1,100-1,200 m ASH (Bostock et al., 2015; Anderson, unpublished data) (Figure 4). Therefore, CWC at Valerie and Forde Guyots were mostly found in water undersaturated in terms of aragonite (Ω_ARAG_ < 1) whilst those at Ghoul and Gothic Seamounts were in water saturated in terms of aragonite (Ω_ARAG_ > 1).

**Figure 4.**
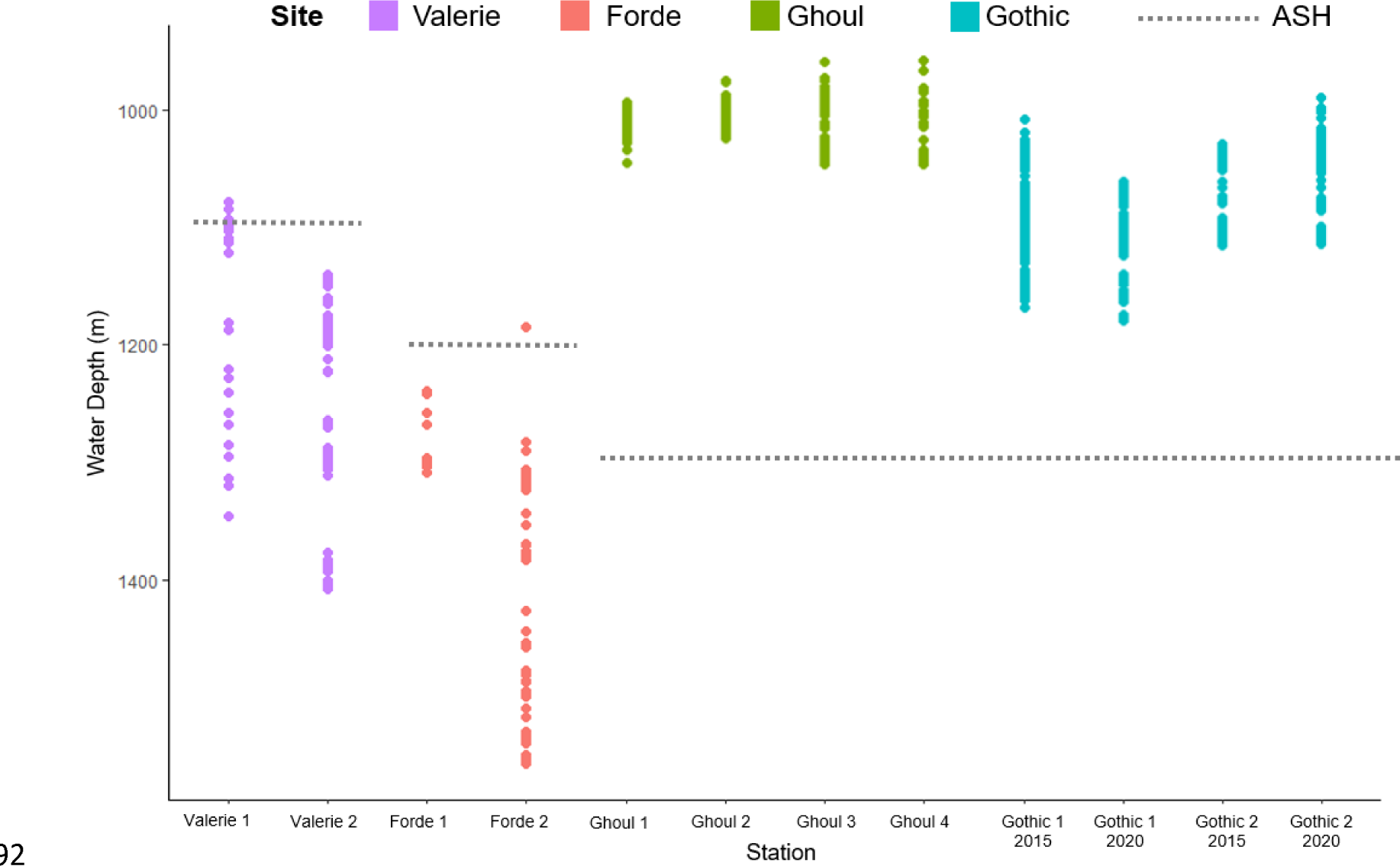
Depth profiles for each station with the average estimated ASH depth for each site. ASH depth for Ghoul and Gothic seamounts was found in Bostock et al. (2015). ASH depths for Valerie and Forde seamounts were provided in a CTD cast summary spreadsheet by Owen Anderson of NIWA.

Across the entire dataset, the mean live:reef ratio was 0.14 (± 0.20). At the Louisville Seamount Chain, the overall mean live:reef ratio was 0.12 (± 0.24), with mean ratios of 0.08 (± 0.18) at Valerie Guyot and 0.18 (± 0.32) at Forde Guyot (Figure 5). At the Graveyard Seamount Complex the overall mean live:reef ratio was 0.15 (± 0.18), with mean ratios of 0.23 (± 0.21) at Ghoul Seamount and 0.12 (± 0.16) at Gothic Seamount (Figure 5).

**Figure 5.**
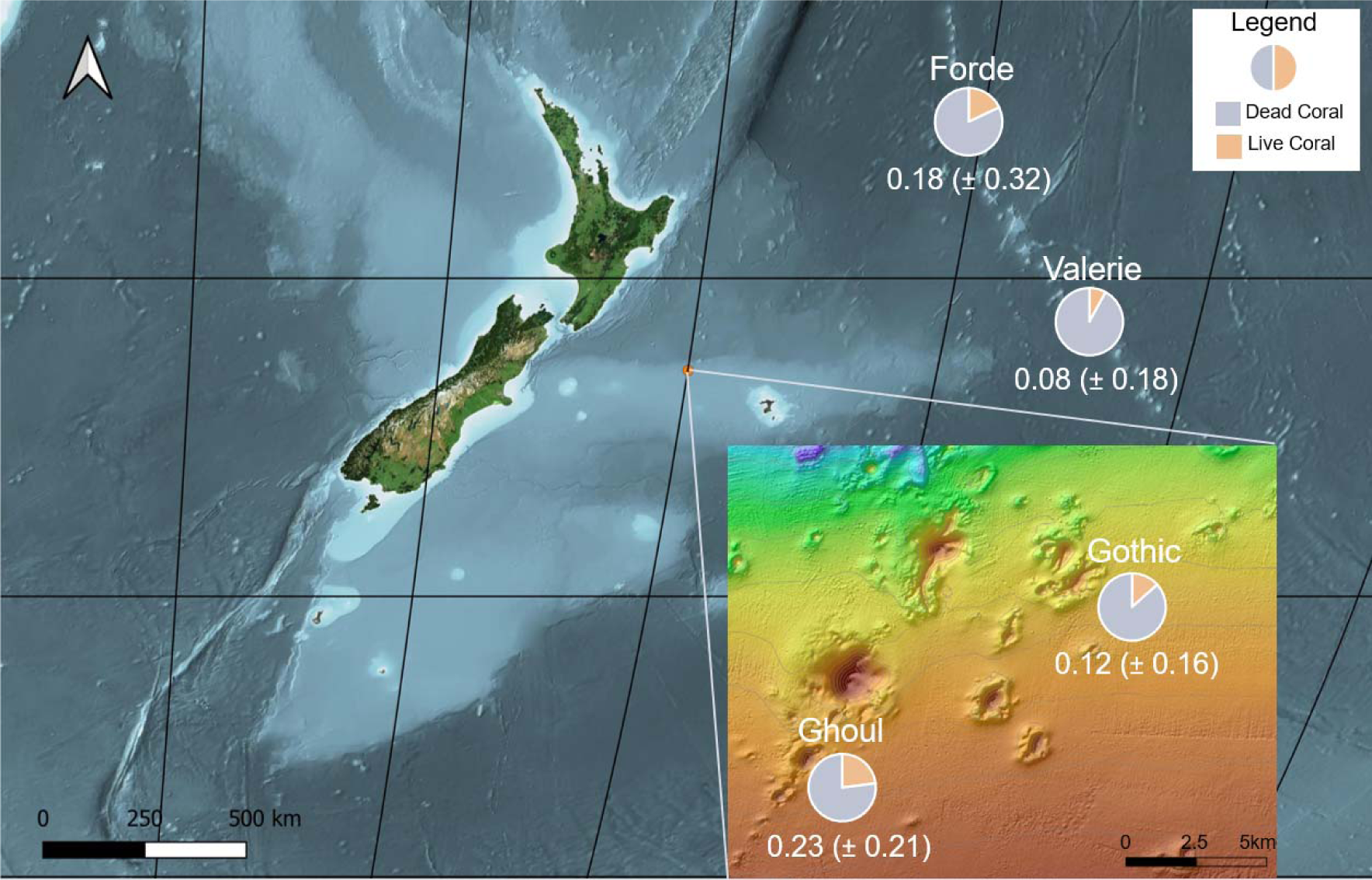
Spatial trends of average (± SD) live:reef ratios across sites.

Significant differences were found for live:reef ratios between stations and sites (p < 0.0001) (Figures 5 & 6). Live:reef ratios were significantly different between the Louisville Seamount Chain and Graveyard Seamount Complex (p < 0.0001). Pairwise comparisons using the Wilcoxon rank sum test showed that most stations live:reef ratios showed statistically significant differences from one another. Stations that did not show statistically significant differences between them were: Valerie 2, Forde 1 and Forde 2; Gothic 1 2015 and Gothic 1 2020; Gothic 2 2015, Gothic 2 2020 and Ghoul 1; Gothic 2 2020 and Ghoul 3; and Ghoul 2, Ghoul 3 and Ghoul 4.

**Figure 6.**
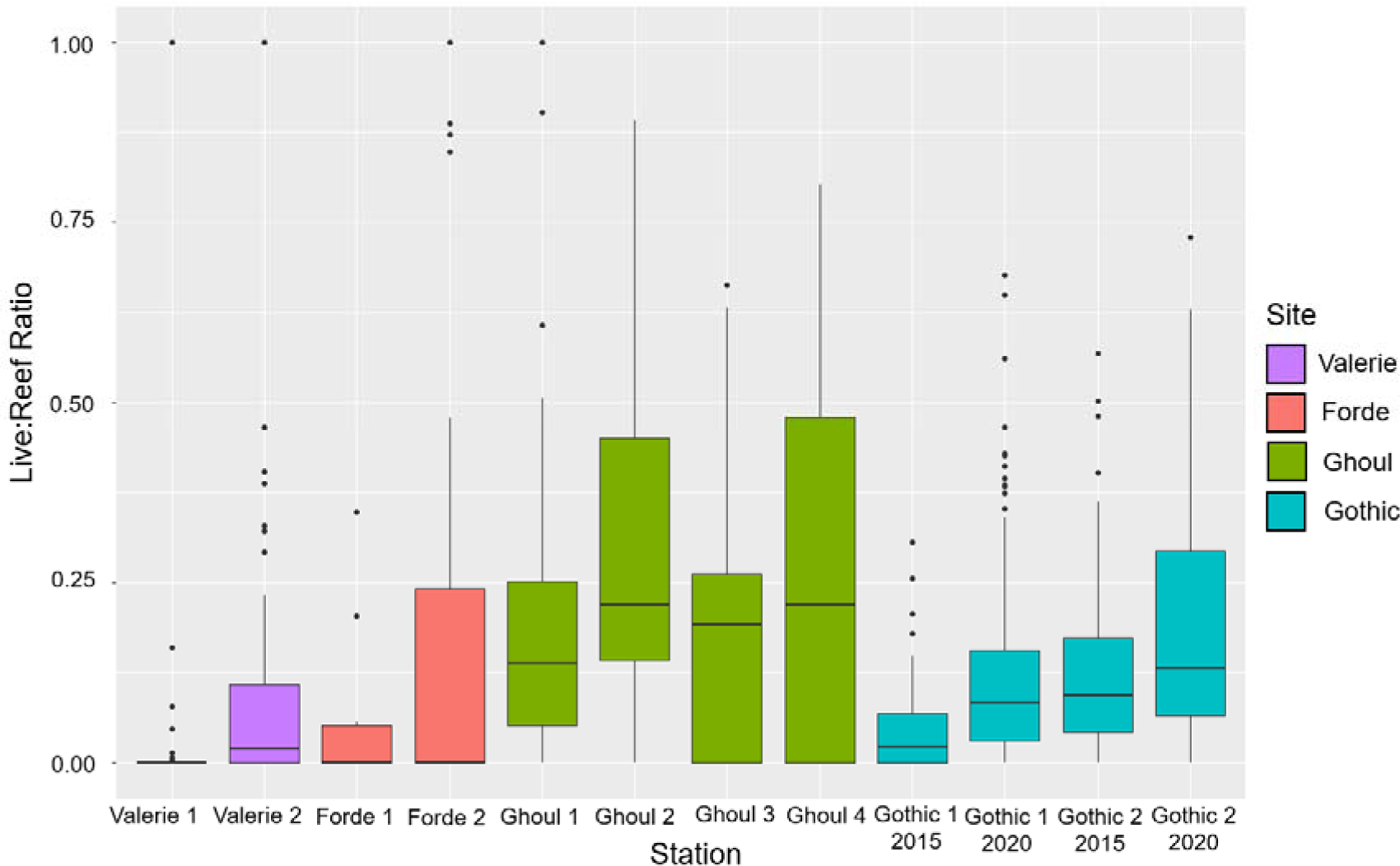
Boxplot of live:reef ratios for each station, grouped by colour based on site. Valerie and Forde sites are situated below the ASH while Ghoul and Gothic sites are situated above the ASH. The bold horizontal line indicates the median and the bounding boxes indicate interquartile ranges.

**Figure 7.**
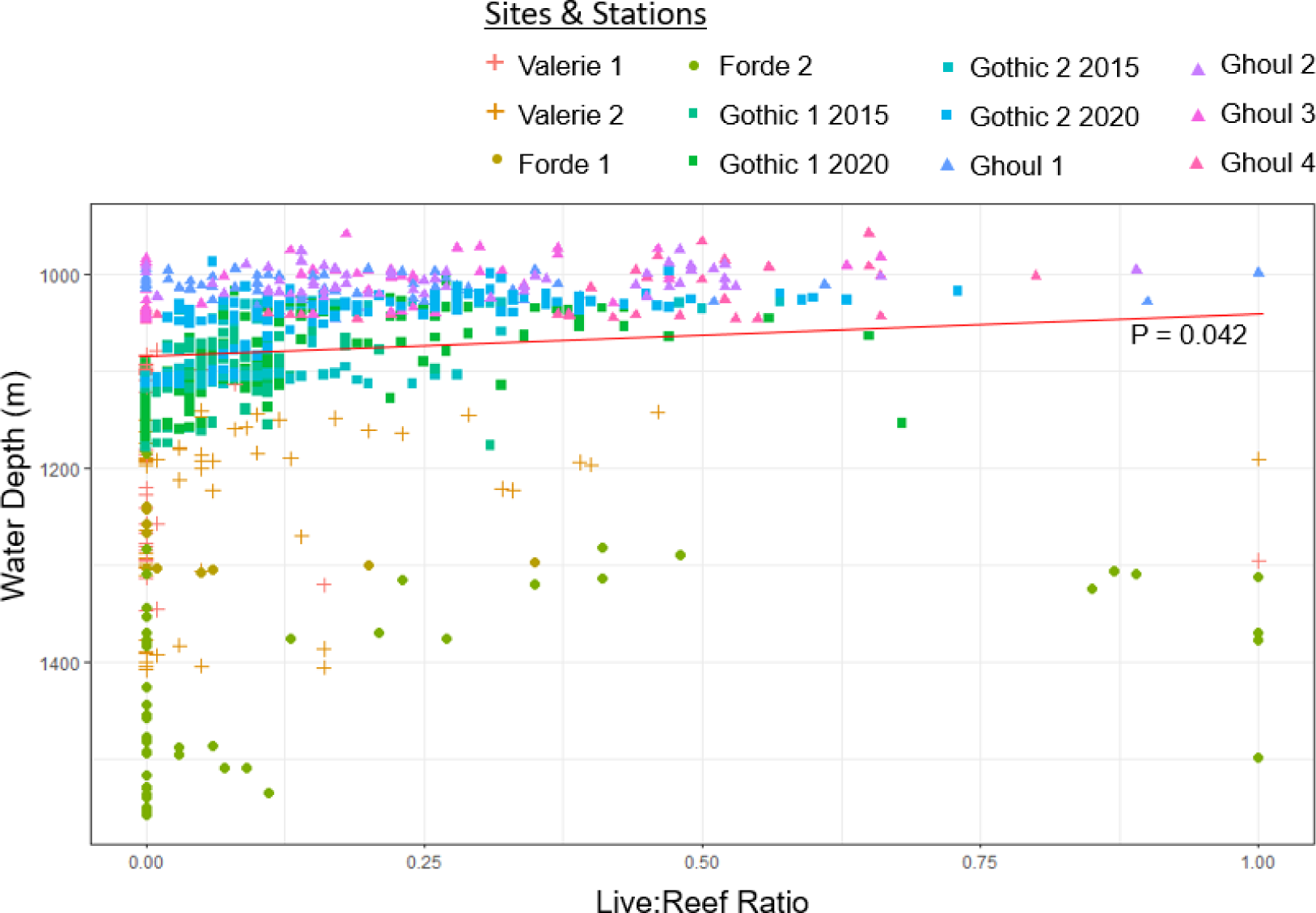
Correlation plot and non-parametric linear model regression between water depth (m) and live:reef ratios for all observations with recorded depth. Different sites are denoted by symbol shape and different stations are denoted by colour and best fit line is shown in red.

The live:reef ratios at Valerie and Forde Guyots within the Louisville Seamount Chain were not significantly different from one another (p = 0.15). Live:reef ratios at the two Valerie Guyot Stations were significantly different from one another (p <0.0001), whereas the two Forde Guyot stations were not (p = 0.62).

The live:reef ratios at the Ghoul and Gothic Seamounts within the Graveyard Seamount Complex were significantly different from one another (p < 0.0001). Live:reef ratios at three of the four Ghoul Seamount stations did not show statistically significant differences from one another (Table 1).

**Table 1.** Pairwise comparisons of live:reef ratios across the four stations at Ghoul Seamount using Wilcoxon rank sum test with continuity correction. Significant differences in live:reef ratios between stations indicated with *.

|  | Ghoul 1 | Ghoul 2 | Ghoul 3 |
| --- | --- | --- | --- |
| Ghoul 2 | 0.006* | - | - |
| Ghoul 3 | 0.029* | 0.541 | - |
| Ghoul 4 | 0.019* | 0.673 | 0.119 |

**Table 2.** Mean surface area of live coral, dead coral, and the entire reef (± standard deviation) for the two Gothic Seamount stations in 2015 and 2020.

| | Live Surface Area in $m^2$ | Dead Surface Area in $m^2$ | Reef Surface Area in $m^2$ |
| --- | --- | --- | --- |
| Gothic 1 2015 | $0.116 \pm 1.01$ | $1.613 \pm 2.53$ | $1.729 \pm 3.20$ |
| Gothic 1 2020 | $0.067 \pm 0.11$ | $0.273 \pm 0.31$ | $0.340 \pm 0.38$ |
| Gothic 2 2015 | $0.464 \pm 1.27$ | $2.464 \pm 2.36$ | $2.928 \pm 3.02$ |
| Gothic 2 2020 | $0.880 \pm 0.10$ | $0.357 \pm 0.22$ | $0.445 \pm 0.26$ |

**Table 3.** Non-parametric linear model results for the relationship between depth and live:reef ratios. * indicates significant values.

| Station | Intercept ± SE | Slope ± SE | p-value | Adjusted R<sup>2</sup> |
| --- | --- | --- | --- | --- |
| Forde 1 | 1306.3 ± 1.6 | -27.1 ± 8.6 | 0.05* | 0.69 |
| Forde 2 | 1437.1 ± 26.6 | -103.3 ± 43.6 | 0.03* | 0.18 |
| Valerie 1 | 1195.7 ± 51.3 | 105.5 ± 133.5 | 0.47 | -0.07 |
| Valerie 2 | 1218.1 ± 18.8 | -46 ± 72.3 | 0.53 | -0.02 |
| Gothic 1 2015 | 1110.99 ± 5.3 | -96.7 ± 60.7 | 0.12 | 0.02 |
| Gothic 2 2015 | 1094.7 ± 4.4 | -162.9 ± 26.1 | <0.001* | 0.27 |
| Gothic 1 2020 | 1098.2 ± 4.2 | -125.2 ± 21.5 | <0.001* | 0.2 |
| Gothic 2 2020 | 1067.8 ± 3.6 | -114.1 ± 14.5 | <0.001* | 0.34 |
| Ghoul 1 | 1009.6 ± 2.5 | 7.4 ± 8.8 | 0.41 | -0.006 |
| Ghoul 2 | 1012.5 ± 2.9 | -18.7 ± 8.2 | 0.027* | 0.07 |
| Ghoul 3 | 1018.5 ± 7.9 | -52.3 ± 26.9 | 0.06 | 0.07 |
| Ghoul 4 | 1051.5 ± 9.6 | -74.1 ± 21.2 | 0.002* | 0.27 |
| Entire Dataset | 1083.3 ± 4.7 | -35.5 ± 17.4 | 0.04* | 0.005 |

## Discussion

As live and dead portions of CWC are threatened by different environmental drivers (Barnhill et al., 2023), this study demonstrates that assessing live:reef ratios could be an important parameter to monitor the long-term health of *S. variabilis* reefs in relation to the ASH. Here we discuss the differences in live:reef ratios across sites and stations and identify potential parameters that could influence the ratios.

### Live:Reef Ratios across CWC Species

This study found higher CWC live:reef ratio occurrences than in previously published studies, likely due to differing methodologies and target species. Vad et al. (2017) and Orejas et al. (2021) have previously reported live:reef ratios of *D. pertusum* and *M. oculata* in the Atlantic Ocean. Vad et al. (2017) did not find any significant differences in live:reef ratios for *D. pertusum* colonies between the onshore Mingulay Reef Complex and offshore PISCES Rockall Bank site in the NE Atlantic despite differences in morphotypes, depth, temperature, salinity, oxygen concentration, and dissolved inorganic carbon.

In the 18 *D. pertusum* colonies analysed by Vad et al. (2017), the live:reef ratio was never higher than 0.27, whereas we found multiple stations with interquartile ranges > 0.25 and some instances with a ratio of 1, where all coral present in an image was living. This means *S. variabilis* analysed in this study has a larger proportion of live coral present than published values for *D. pertusum*. However, this study did use a different method than Vad et al. (2017), where ratios were quantified through linear extension within colonies as opposed to the semi-automatic method of measuring surface area. Orejas et al. (2021) found larger live:reef ratios than Vad et al. (2017), though still lower than those found in this study, with an average of 0.36 and a maximum of 0.54. Tracey et al. (2013) observed that like its Atlantic counterparts, the living portion of *S. variabilis* is significantly smaller than the dead supporting skeletal framework.

### Live:Reef Ratios between South Pacific Ocean Sites

Live:reef ratios between stations were significantly different from one another at Valerie Guyot but not at Forde Guyot. These discrepancies between features could be explained by inter-feature variability in environmental conditions, such as current speed and food supply. At a single seamount, local topography and location on the feature can impact waterflow and with it, food supply (Genin et al., 1986), which may have drove the differences seen in this study. CWC are often found in areas with strong currents, as they provide higher levels of food (Lim et al., 2020; Mohn et al., 2014). The two stations analysed at Valerie Guyot were from different flanks ∼15 km apart, while the two at Forde were adjacent to one another (< 0.5 km apart), indicating more similar environmental conditions for the two stations at Forde Guyot than the two stations at Valerie Guyot. Sites near each other showed more similarities in live:reef ratios than sites further apart from one another, suggesting similar environmental conditions result in similar live:reef ratios.

Framework forming corals at Gothic Seamount have a mean percentage cover of 21% (Clark & Rowden, 2009). In addition to *S. variabilis*, the framework forming coral *Madrepora oculata* is also present at Gothic Seamount (Clark & Rowden, 2009). The live:reef ratios at Gothic Seamount remained statistically indistinguishable between 2015 and 2020 despite significant differences in the total area of live coral, dead coral, and the entire reef present in analysed images. This suggests the environmental parameters driving ratios remained constant over time at this site. The variability observed in reef size parameters was likely due to some differences in the actual DTIS tracks over the seamount. Although the survey design tried to replicate transects as closely as possible, there was slight variability in the DTIS track positions given weather and near-seabed current conditions (Clark et al. 2019).

While uncommon, surveys have observed trawl gear impacts at Gothic Seamount (Clark & O’Driscoll, 2003; Clark & Rowden, 2009). The largely untrawled Ghoul and Gothic Seamounts have higher CWC cover than other features in the Graveyard Seamount Complex due to heavy trawling in surrounding areas (Clark et al., 2019); Therefore, live:reef findings for these sites should not be extrapolated for nearby features where trawling activity may have impacted live:reef ratios.

We speculate that different environmental conditions and colony genetics between the Louisville Seamount Chain and Graveyard Seamount Complex drove the statistically significant differences in live:reef ratios between sites. The ranges of depth, temperature, and aragonite saturation were different between the two areas. Additionally, subtropical surface waters and subantarctic waters from the Southern Ocean meet near the Graveyard Seamount Complex, causing higher primary productivity than is found at the Louisville Seamount Chain (Tracey et al., 2023). Higher than average surface productivity provides greater levels of food to CWC, driving reef distribution for several species, including *S. variabilis* (Maier et al., 2023). The corals in this study are also genetically different between the two regions (Zeng et al., 2017), which could explain some of the observed differences in live:reef ratios. Distance between *S. variabilis* colonies has a significant impact on microsatellite genetic isolation and individuals in the southern region (including the Graveyard Seamounts) have greater nucleotide and haplotypic diversity than those in the Northern region (including the Louisville Seamount Chain) indicating differences in population structure (Zeng et al., 2017).

### Potential Environmental Parameters Impacting Live:Reef Ratios

This study found depth to be a driver of live:reef ratios across the entire dataset. Depth acts as a main predictor for *S. variabilis* distribution in New Zealand waters and the Louisville Seamount Chain (Rowden et al., 2017; Tracey, Rowden, et al., 2011), and similarly, was found to contribute to live:reef ratios in this region. Across this data set, the general trend saw greater live:reef ratios at shallow depths, indicating there is more live coral at shallow depths and more dead coral at deeper depths. However, this result is not consistent across all stations as p-values did not indicate significant results at five of the twelve stations, indicating there are additional drivers of live:reef ratios. Interestingly, Ghoul 2 and Ghoul 4 have the widest spread of ratios despite a relatively small depth range compared to other stations. More work is needed to identify additional drivers of variability in live:reef ratios, in order to better predict reef habitat futures at different sites under climate change. Finer-scale measurements of environmental parameters are required to identify other drivers of live:reef ratios for *S. variabilis*.

Compared to preindustrial levels, the ASH across the New Zealand exclusive economic zone has already risen by 50-100 m (Mikaloff-Fletcher et al., 2017). Regional environmental and spatial variability cause ASH depth variations at Chatham Rise, where the 1,300 m ASH at the Graveyard Seamount Complex occurs 300 m deeper than the 1,000 m ASH south of the rise (Bostock et al., 2015). Differences in aragonite saturation on the features are caused by patterns of oceanic circulation whereby certain water masses are physically blocked by complex topographies, such that the older and more corrosive North Pacific Deep Water influences bottom waters north of Chatham Rise (Law et al., 2018; Tracey et al., 2013).

Despite New Zealand CWC occurrences below the ASH, Thresher et al. (2011) concluded that aragonite saturation state likely constrains colonial CWC taxa distributions, whereas it has less effect on solitary coral species. Additional research suggests CWC living below the ASH likely became established when the ASH was deeper, and have since adapted to the undersaturated conditions (Mikaloff-Fletcher et al., 2017). Using the 2100 projected ASH depth of 600 m, only Chatham Rise, parts of the Campbell Plateau, some Kermadec and Macquarie Ridge seamounts, and the New Zealand continental shelf will remain in aragonite saturated waters (Bostock et al., 2015). Gammon et al. (2018) found reduced coenenchyme tissue coverage and elevated respiration rates in *S. variabilis* when exposed to water undersaturated in aragonite, indicating the shoaling ASH and lower pH could impact physiological responses from the live coral colonies as well as cause dead intact framework dissolution. From the live:reef results of this study, we can estimate the amount of dead intact framework threatened by the shoaling ASH at each site (ranging from 77% at-risk reef surface area at Ghoul Seamount to 92% at-risk reef surface area at Valerie Guyot), which can help inform which sites should be selected for protection as possible climate change refugia. We can also infer the shoaling ASH will first impact reef areas with more dead intact framework, as live:reef ratios are lower at deeper depths.

Additional evidence has shown that conditions below the ASH are not optimal where dead intact framework has been documented covered in metal oxide coating, preserving the 3-dimensional reef matrix, but without the complete biological associates, suggesting it is not an ideal habitat for benthic organisms (Thresher et al., 2011). Dead intact framework of *S. variabilis* reefs coated in ferromanganese oxide crust have been documented in the southwest Pacific at both fished and unfished seamounts located in the Huon Marine Reserve, Tasman Fracture Zones, and Louisville Seamount Chain (Clark et al., 2014; Thresher et al., 2011). This phenomenon was also seen in this study at the Louisville Seamount Chain. Predicted to be over 1,000 years old (Clark et al., 2014), this dead intact framework is preserved in the ferromanganese oxide crust at Forde Guyot despite existing beneath the ASH. While this preserves the framework’s 3-dimensional matrix, the structure is no longer a functional reef, as the loss of living coral leads to loss of ability to provide key ecosystem services to support biodiversity.

## Conclusion

This study explored live:reef ratios in relation with the depth, as a proxy for ASH, temperature, for the framework-forming species *S. variabilis* in the southwest Pacific Ocean. Across the dataset, depth was found to be a driver of live:reef ratios, with higher live:reef ratios at shallower depths. Through image analysis, we found significant differences in live:reef ratios between the Louisville Seamount Chain and the Graveyard Seamount Complex. Along with depth, we speculate that environmental differences in temperature and Ω_ARAG_ combined with genetic variability may explain these differences. We found live:reef ratios remained constant between 2015 and 2020 at the Gothic Seamount despite significant changes in surface area of live corals and dead intact framework of *S. variabilis* as well as the entire reef structure. While more data is needed to establish if Ω_ARAG_ and genetic variability indeed contributed to the ratio differences, the conducted dead framework measurements may indicate the amount of dead intact framework CWC threatened by the shoaling ASH at each site, which could help inform which sites should be selected for protection as possible climate change refugia.

## Competing Interests

The Authors declare no competing interests.

## Acknowledgements

The authors would like to thank NIWA scientists Owen Anderson for providing CTD data and Susi Woelz for providing bathymetry grids for this research. The videos from TAN1402 were collected by NIWA as part of the South Pacific Vulnerable Marine Ecosystems Project (C01X1229) funded by Ministry of Business, Innovation & Employment. The videos from TAN1503 were collected by NIWA as part of the “Impact of resource use on vulnerable deep-sea communities” project (CO1X0906), funded by the Ministry of Business, Innovation & Employment with support from Ministry for Primary Industries (project BEN2014-02) and NIWA’s Enabling the Management of Marine Mining (MBIE contract CO1X1228). The videos from TAN2009 were collected by NIWA as part of the “Recovery of seamount communities” project (ZBD2020-07), funded by the Ministry for Primary Industries, with vessel time provided by the Ministry for Business, Innovation, and Employment. This work was supported by the Independent Research Fellowship from the Natural Environment Research Council (NERC) to SJH (NE/K009028/1 and NE/K009028/2)

## Author Contributions

**Kelsey Archer Barnhill**: Conceptualization, Methodology, Formal Analysis, Writing – Original Draft, Visualization. **Caroline Chin**: Resources, Data Curation, Writing – Review & Editing, Project Administration. **Di Tracey:** Resources, Writing – Review & Editing, Project administration. **Malcolm Clark**: Resources, Writing – Review & Editing, Funding Acquisition**. Laurence H. De Clippele:** Methodology, Software, Writing – Review & Editing. **Sian F. Henley**: Writing – Review & Editing, Supervision. **Uwe Wolfram:** Conceptualization, Writing – Review & Editing, Supervision. **Sebastian Hennige**: Conceptualization, Resources, Writing – Review & Editing, Supervision.

## Notes

### Competing Interest Statement

The authors have declared no competing interest.

